# Motivated for near impossibility: How task type and reward modulate task enjoyment and the striatal activation for extremely difficult task

**DOI:** 10.1101/828756

**Authors:** Michiko Sakaki, Stef Meliss, Kou Murayama, Yukihito Yomogida, Kaosu Matsumori, Ayaka Sugiura, Madoka Matsumoto, Kenji Matsumoto

## Abstract

Economic and decision-making theories suppose that people would disengage from a task with near zero success probability, because this implicates little normative utility values. However, humans are often motivated for an extremely challenging task, even without any extrinsic incentives. The current study aimed to address the nature of this challenge-based motivation and its neural correlates. We found that, when participants played a skill-based task without extrinsic incentives, their task enjoyment increased as the chance of success *decreased*, even if the task was almost impossible to achieve. However, such challenge-based motivation was not observed when participants were rewarded for the task or the reward was determined in a probabilistic manner. The activation in the ventral striatum/pallidum tracked the pattern of task enjoyment. These results suggest that people are intrinsically motivated to challenge a nearly impossible task, but only when the task requires certain skills and extrinsic rewards are unavailable.

## Introduction

In December 2015, the Government Communications Headquarters in the United Kingdom released a Christmas puzzle. The puzzle was allegedly extremely difficult and time consuming at the time of announcement, and apparently promised no monetary rewards for solving it. Nevertheless, more than 600,000 people tried to solve the puzzle and (unfortunately) no one succeeded. As illustrated in the example, humans seem to have a natural inclination to challenge an extremely improbable outcome, even without extrinsic incentives (e.g., monetary rewards). It is difficult to explain such challenge-oriented motivation (Loewenstein, 1999), because there is little chance of succeeding in the task, making normative utility value almost zero according to standard economic, decision-making, and reinforcement-learning theories.

The purpose of the current article is to examine the conditions under which people express such motivation for nearly impossible outcomes and its neural correlates. The dopaminergic reward network is activated more strongly when people are presented with cues signalling higher expected reward value (Bartra et al., 2013; Delgado, 2007; Diekhof et al., 2012). Based on these findings, the reward network is expected *not* to be activated by cues that predict a task with extremely-low chance of success. Past studies on reward and decision-making, however, overlooked two critical factors for understanding the task motivation for nearly impossible outcomes. First, previous studies mainly employed tasks in which outcome was decided in a pure probabilistic manner (luck task). In contrast, we are often motivated for challenging tasks in our daily life when the task requires some skills. When task success is contingent upon one’s skills, people focus more on the improvement of their skills, rather than task success itself; as a result, people may exhibit greater persistence after failure (Lee & Kim, 2014; Lee & Reeve, 2017), perhaps even when the task is almost impossible to achieve.

Second, previous research has typically implemented extrinsic rewards to quantify the value of task cues. However, humans are often intrinsically motivated for a challenging task in the absence of extrinsic incentives (Csikszentmihalyi, 1990; Deci & Ryan, 1985; Murayama, 2022). In fact, performance-contingent extrinsic rewards can put people in a conflicting situation in the face of a challenging task. Specifically, people’s motivation for challenge may be offset by the risk of losing out potential extrinsic rewards. In other words, there may be a qualitative difference in people’s motivation when extrinsic reward is at stake and when it is not (Baranes, 2014; Braver et al., 2014; Murayama et al., 2010; Ten et al., 2021).

We examined whether the nature of the task and the presence of extrinsic rewards influence people’s motivation for nearly impossible outcomes and how these factors influence neural responses to the task with different chance of success (including a near impossible task). Participants were engaged in a game-like task (Figure 1A) in which participants were asked to press a button to stop a stopwatch. A trial is regarded as success if participants were able to stop the stopwatch within a specific time window, which is around 5 seconds after the stopwatch starts. Some participants performed this skill task without performance-contingent extrinsic rewards (no-reward group), while others played the task with outcome-contingent monetary rewards (reward group). The remaining participants (gambling group) played a similar task, but they simply received extrinsic rewards that were determined in a pure probabilistic manner (i.e. luck task). More specifically, they were simply asked to watch the stopwatch starting and stopping on its own. If the stopwatch stopped within the time window, participants earned monetary rewards. Across all groups, each trial was preceded by a cue indicating the success probability of the upcoming task --- high-, moderate-, and extremely-low chance (of success) conditions.

**Figure 1.**
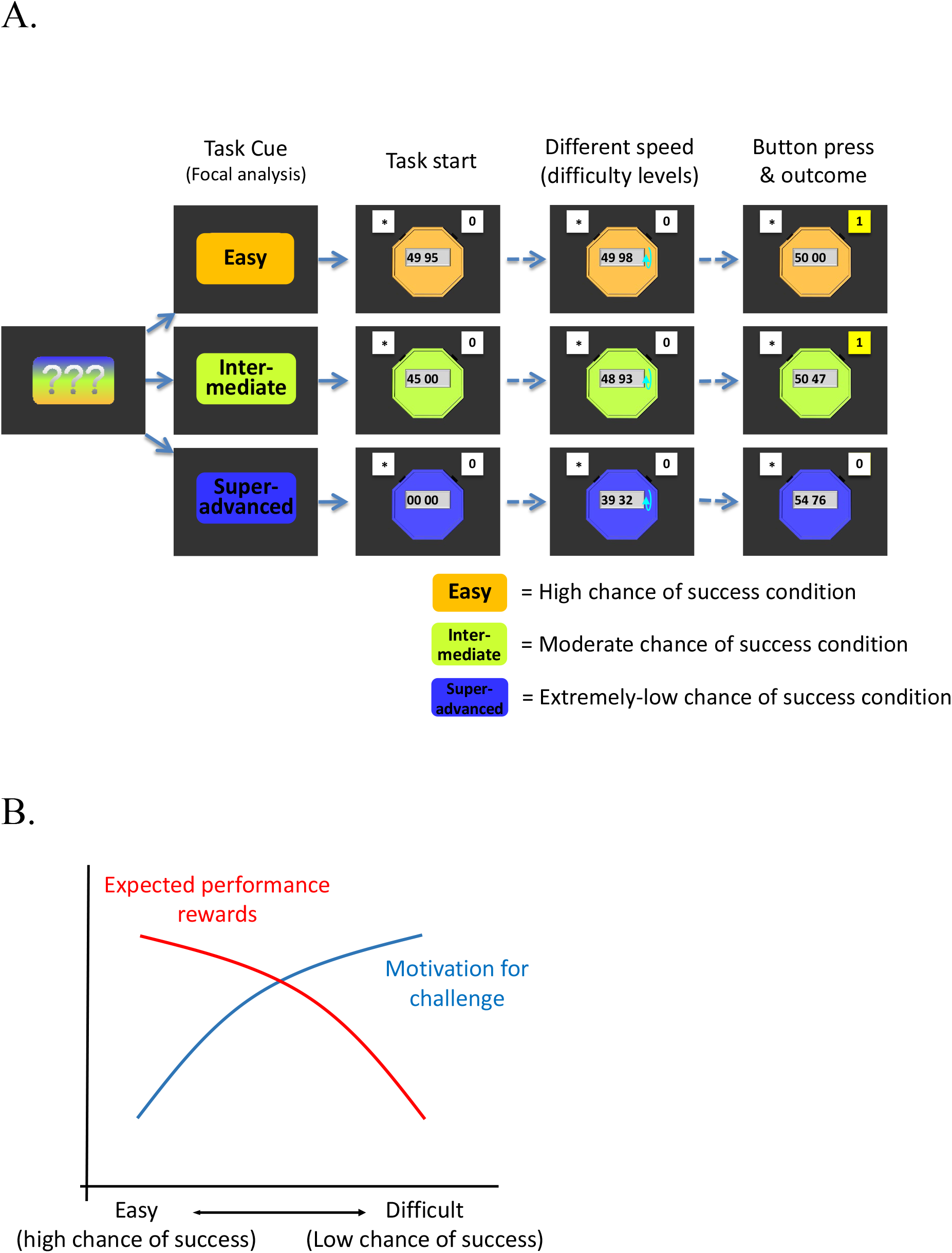
**A**. Task sequence. Participants were presented with a task cue signalling different chance of success (“Easy” = high chance of success, “Intermediate” = moderate chance of success, and “Super-advanced” is extremely low chance of success), followed by the actual task. There were also control trials (“watch-stop” trials) in which participants neither succeeded nor failed, but these trials were omitted from the figure. **B**. A schematic picture of how expected performance rewards (red line) and motivation for challenge (blue line) changes in an opposite manner as a function of task difficulty (chance of success). Expected performance reward decreases as chance of success decreases (i.e. low expected utility), whereas motivation for challenge *increases* as chance of success decreases.

We expected that participants would exhibit increasing overall motivation for the task with lower chance of success only when they were engaged in a skill task without extrinsic incentives (no-reward group; Figure 1B, blue line). As such, we predicted that participants’ motivation would monotonically *increase* as the chance of success decreases, even down to the point when the task was almost impossible to achieve. On the other hand, when participants were engaged in a luck task with extrinsic incentives (gambling group), there was no room for people to focus on the improvement of their skills. As such, we predicted that participants’ motivation would monotonically *decrease* as the chance of winning rewards decreased, simply because a task with higher chance of success involves higher expected performance reward (Figure 1B, red line). When participants were engaged in a skill task with extrinsic rewards (reward group), participants’ motivation to challenge improbable outcome and motivation to obtain high extrinsic rewards contradict each other (Figure 1B, blue line and red line). As a result, we did not have a strong prediction: we rather expected relatively unclear or weak relationship between the chance of task success and motivation.

To assess motivation, we asked about task enjoyment for the task with different difficulty levels (i.e. the extent to which participants enjoyed the respective task). Task enjoyment represents the subjective positive emotional experience during the task, which should reflect the downstream process of motivational engagement. Thus, the measure should be suitable to gauge overall motivation incorporating different sources of motivation (i.e. motivation for extrinsic rewards and motivation for challenge; Figure 1B). To investigate how challenge-based motivation is represented in the brain, we also examined the brain activation in response to the task cue signalling the difficulty level (i.e. anticipatory phase of the task). Reward-related areas are obvious candidate areas (the striatum, the ventromedial prefrontal cortex, midbrain, etc.), and among these regions, previous studies using the same task showed that the striatal activation seems to most clearly reflect motivation for the task (Murayama et al., 2010, 2015). Thus, our neuroimaging analysis primarily focused on the cue-related activation in the striatum as an a priori region of interest.

## Materials and Methods

### Participants

Fifty-five participants (mean age = 20.05, *SD* = 1.37; 28 males) were allocated to the no-reward group, the reward group, or the gambling group, constituting a 3 (group, between subjects: no-reward, reward, or gambling) x 3 (chance of success, within-subjects: high chance, moderate chance, or extremely-low chance) factorial design. Participants in the reward and no-reward groups were randomly assigned to these groups, and the data for the gambling group were later collected. Prior to data analyses, we excluded data from four participants: one due to a technical problem, another due to an incidental finding (i.e. medical abnormality), one due to large motions, and the other because the participant failed to press buttons in most trials. This resulted in 17 participants in the no-reward condition, 17 participants in the reward condition, and 17 in the gamble condition. Thus, the final study sample consisted of *N*= 51 adults aged between 18 and 24 (*M* = 20.1, *SD* = 1.39), with 23 males and 28 females.

To determine the sample size, we simply collected as much data as possible within a specific time window. Our previous study (Murayama et al., 2010) found that the effects of reward context had a large effect on the striatal activation (the region of interest in the current study) in a game-like task that we used (*N* = 28, between-subjects, two group design). Accordingly, we considered that the sample size is sufficient to detect the effects of reward context manipulation.

### Experimental Procedures

Participants played a stopwatch task, which was designed based on the task used in previous research (Murayama et al., 2010, 2015). Participants’ task was to stop a stopwatch within a specific time window --- between 49.95 and 50.05. There were three types of stopwatches which are different in terms of the chance of success: a) in the high-chance condition, the stopwatch started from 49.95 and ran 100-time slower than an ordinary watch; b) in the moderate-chance condition, the stopwatch started from 45.00 and ran in the same speed as an ordinary watch (like the ones used in previous research); c) in the extremely-low chance condition, the stopwatch started from 0.00 and ran 10-time faster than an ordinary watch. Thus, while the initial time was different across the three types of the stopwatches, they were different in their speed, allowing us to manipulate task difficulties while making the trial duration similar across the conditions (e.g., in all conditions participants needed to press a button around 5 seconds after the stopwatch started in order to stop the stopwatch at 50.00).

In the no-reward and the reward groups, each trial (Figure 1) started with three question marks. Upon participants’ button press, the question marks were replaced by a word cue indicating the task difficulty level (i.e. chance of success) of the subsequent stopwatch: “easy” (i.e. high chance of success), “intermediate” (i.e., moderate chance of success), and “super advanced” (i.e. extremely-low chance of success). The cues (and the following stopwatches) were differently colored depending on the chance of success (blue, green, and orange), and the assignment of the colors was counterbalanced across participants. Participants were then shown the stopwatch, and asked to press a button with the right thumb to stop it right between 49.95 and 50.05. The stimulus onset asynchrony (SOA) of the cue presentation and start of the stopwatch task was jittered between 4.5 and 8.5 seconds. Immediately after they pressed the button, they were shown success feedback or failure feedback for 0.8 seconds, followed by a jittered ITI between 3 and 7 seconds. This task sequence was optimized to examine the brain activation associated with the cue presentation. Unbeknownst to participants, in the extremely-low chance condition, the displayed time when they stopped the stopwatch was manipulated so that participants always failed. In the moderate chance condition, we made an online adaptive adjustment of task difficulty based on participants’ previous task performance so that most participants achieved approximately 50% accuracy. Specifically, the algorithm calculated the overall success rate in the previous trials for the moderate chance condition, and made the task slightly more difficult when the overall success rate was more than 50%, whereas the task was made slightly easier when the overall success rate was less than 50%. If the success rate was 50%, no change was made. The adjustment of difficulty level was achieved by secretly increasing or decreasing the actual time window for success. In the high-chance condition, the task was so easy that participants succeeded in all trials without any manipulations. No participants indicated the possibility that their task performance had been manipulated after the experiment. If participants pressed a button too early or too late (e.g., if participants did not press a button within 10 secs after the stopwatch started), a warning message “please press a button” appeared and this trial was skipped.

Prior to the session, participants were told about the differences of the three different types of stopwatches. Specifically, participants were told that the stopwatch game in the high chance condition would be very easy and they can easily win; that the moderate chance condition would be moderately difficult; and that the extremely-low chance condition would be extremely difficult and would be almost impossible to win. Participants in the reward group were further told that they would earn 100 yen (approximately USD 1) for every success irrespective of the type of stopwatch. Participants in the no-reward group were not told anything about performance-based monetary rewards.

The procedure in the gambling group was similar to that in the reward group with a few exceptions. As in the reward group, participants were told that they would earn 100 yen for each time that the stopwatch stopped between 49.95 and 50.05. However, unlike the reward group, they did not have control over the timing at which the stopwatch stopped and the stopwatch was called a lottery machine. They were simply asked to watch the stopwatch starting and stopping on its own and press the button after the stopwatch stopped. If the stopwatch stopped within the time window, this trial was regarded as hit (success) and participants earned money. Participants were told that the outcome would be determined by the computer and the task cue would indicate the chance of hit (success) in that trial. Specifically, they were instructed that it would be almost certain that they can earn money in the high-chance condition, whereas it would be almost impossible that they would earn money in the extremely-low chance condition. They were also told that there would be a fair chance (50%) of earning money in the moderate chance condition.

In all three groups, participants also completed a control condition, in which they simply viewed a stopwatch starting and stopping on its own and pressed the button after the stopwatch stopped without any monetary rewards (watch-stop condition). The watch-stop condition was preceded by a word cue “passive viewing”, and the color of the cue and the stopwatch was always gray. The stopwatch ran in one of three different speeds (randomly chosen), which were matched to one of the three chance-of-success conditions). If participants did not press a button within 2 seconds after the stopwatch stopped, a warning message “please press a button” appeared and this trial was repeated.

Across all groups, participants completed 80 trials (20 trials with high chance of success, 20 trials with moderate chance of success, 20 trials with extremely-low chance of success, and 20 watch-stop trials) that were divided into two runs of 40 trials. The sequence of the trials (i.e. the trial order of conditions) was randomized across participants, but to control for any potential trial order effects, we used the same set of trial sequences across the three groups. In addition, the sequence of success and failure trials in the gambling group was matched to that in the reward group. After the session, they completed a self-reported questionnaire (on a 1-7 scale) asking about their task enjoyment to engage in the three types of stopwatch tasks (three items taken from a previous study; Elliot & Harackiewicz, 1996; e.g., “It was enjoyable to play the stopwatch of high chance of success”, “It was boring to play the stopwatch of high chance of success” [reverse coded], and “Stopwatch of high chance of success was fun”). We used exactly the same wording for all the groups. Separate from these questions, participants also answered an exploratory measure assessing participants’ emotional feeling about the task cue (not the task itself) --- anticipated emotional value. The measure consists two questions about their perceived emotional value in response to the task cue for each of the three stopwatches (“The cue for the moderately difficult stopwatch made me pleased”, “The cue for the extremely difficult stopwatch made me disappointed”, reverse coded, *r*s = .67 - 89).

### fMRI Acquisition and Preprocessing

The functional imaging was performed on a 3T magnetic resonance imaging (MRI) scanner (MAGNETOM Trio, A Tim System, Siemens, Germany) with a 32-channel matrix head coil at the Tamagawa University with gradient echo T2*-weighted echo-planar imaging (EPI) sequence. The imaging parameters were TR = 2500 ms, TE = 25 ms, slice thickness = 3 mm, and flip angle = 90°. The data were preprocessed using Statistical Parametric Mapping 12 (SPM12, http://www.fil.ion.ucl.ac.uk/spm/software/spm12/). Images were motion-corrected with realignment to the first volume of the session, corrected for slice-timing, co-registered to the bias-corrected, segmented structural image, spatially normalised to the standard Montreal Neurological Institute (MNI) EPI, and spatially smoothed using a Gaussian kernel with a full width at half-maximum (FWHM) of 8 mm.

### fMRI Data Analysis

For each participant, the blood oxygen level-dependent (BOLD) response was modeled with the general linear model (GLM) for the following regressors of interest: the high-chance stopwatch cue, the moderate-chance stopwatch cue, the extremely-low-chance stopwatch cue, and the three watch-stop cues (i.e. watch-stop cues with three different speeds; they were separately modeled and then averaged). In addition, feedback for stopwatch trials, error trials (i.e. participants did not press a button for a certain duration; see experimental procedures; for these trials, the entire duration of the trial was modeled), session effects, and motion parameters were included as regressors of no interest. The regressors (except motion parameters and session effects) were calculated using a boxcar function for each stimulus convolved with a canonical hemodynamic response function (HRF) without derivatives. Temporal autocorrelation was accounted for by using a first-order autoregressive model during Classical (ReML) parameter estimation. Results for the three watch-stop cues were averaged and used in the following three contrasts to examine the effects of different cues: (i) a contrast between the high-chance stopwatch cue and the averaged watch-stop cues; (ii) another contrast between the moderate-chance stopwatch cue and the averaged watch-stop cues; and (iii) a contrast between the extremely-low chance stopwatch cue and the averaged watch-stop cues.

The resulting contrast images were submitted to a 3 (chance of success: high chance, moderate chance, or extremely-low chance) x 3 (group: no-reward, reward, or gambling) mixed ANOVA with chance of success as within-subject factor and group as between-subject factor. Our primary analysis focused on the reward network in the brain, especially the striatum/pallidum. Thus, we performed a region of interest (ROI) analysis. The striatum/pallidum ROI mask included the bilateral caudate, putamen, and pallidum from the Automated Anatomical Labeling (AAL) atlas (Tzourio-Mazoyer et al., 2002). For the purpose of completeness, we also conducted a set of whole-brain GLM analyses. We applied a voxel-level family-wise error (FWE)-corrected threshold (*P_FWE_* < 0.05, *k* ≥ 5) for ROI analysis as well as the whole-brain analysis.

### Functional Connectivity Analysis

We also conducted a functional connectivity analysis, i.e., generalised psychophysiological interaction (gPPI; McLaren et al., 2012), in order to examine which brain regions were co-activating with the ventral striatum/ventral pallidum during the cue presentation. To perform this analysis, a separate GLM was constructed in which the time course extracted from a seed region (“physiological main effect”) is multiplied with the task time course (“psychological main effect”) to derive with a psychophysiological interaction term that is used as predictor within the GLM. Thereby, gPPI allows us to identify areas with time courses that are better predicted by the time course of the seed region in one psychological context (i.e., task condition) than in others. To model the task time course, the onset and duration of the cue images for stopwatch and watch-stop task at each chance of success were used.We were interested in the difference between the stopwatch and watch-stop cue at each level of chance of success. The analyses were performed using the gPPI toolbox, and the ROI masks were created using MarsBar (Brett et al., 2002). We used a cluster-extent FWE-corrected threshold (FWE < 0.05) with the cluster-defining threshold of *p* = 0.005, to which Bonferroni correction for two separate seed regions (i.e., the left and right ventral striatum/ventral pallidum) was applied, resulting in a cluster-defining threshold of *p* = 0.0025.

## Results

### Behavioural results

The ratings of the task enjoyment were analysed using a 3 (group: no-reward, reward, or gambling, between-subjects) x 3 (chance of success: high chance, moderate chance, or extremely-low chance, within-subjects) mixed ANOVA. This ANOVA showed a main effect of chance of success, *F*(2,96) = 10.02, *P* < .001, generalized η^2^ = 0.13, which was further qualified by the interaction between group and chance of success, *F*(4,96) = 22.72, *P* < .001, generalized η^2^ = 0.41. The main effect of group was not statistically significant, *F*(2, 48) = 1.90. The pattern of the results confirmed our prediction (Figure 2A). Specifically, participants in the no-reward group showed a linear pattern with an *increase* in task enjoyment when the chance of success decreased, exhibiting greatest task enjoyment for improbable outcomes. On the other hand, participants’ task enjoyment in the gambling group *decreased* as the chance of success decreased, showing an opposite linear pattern from that in the no-reward group. Finally, in the reward condition, task enjoyment was numerically highest in the moderate chance condition (inverted-U trend), but the pattern does not seem to be clear enough in comparison to the other two groups.

**Figure 2.**
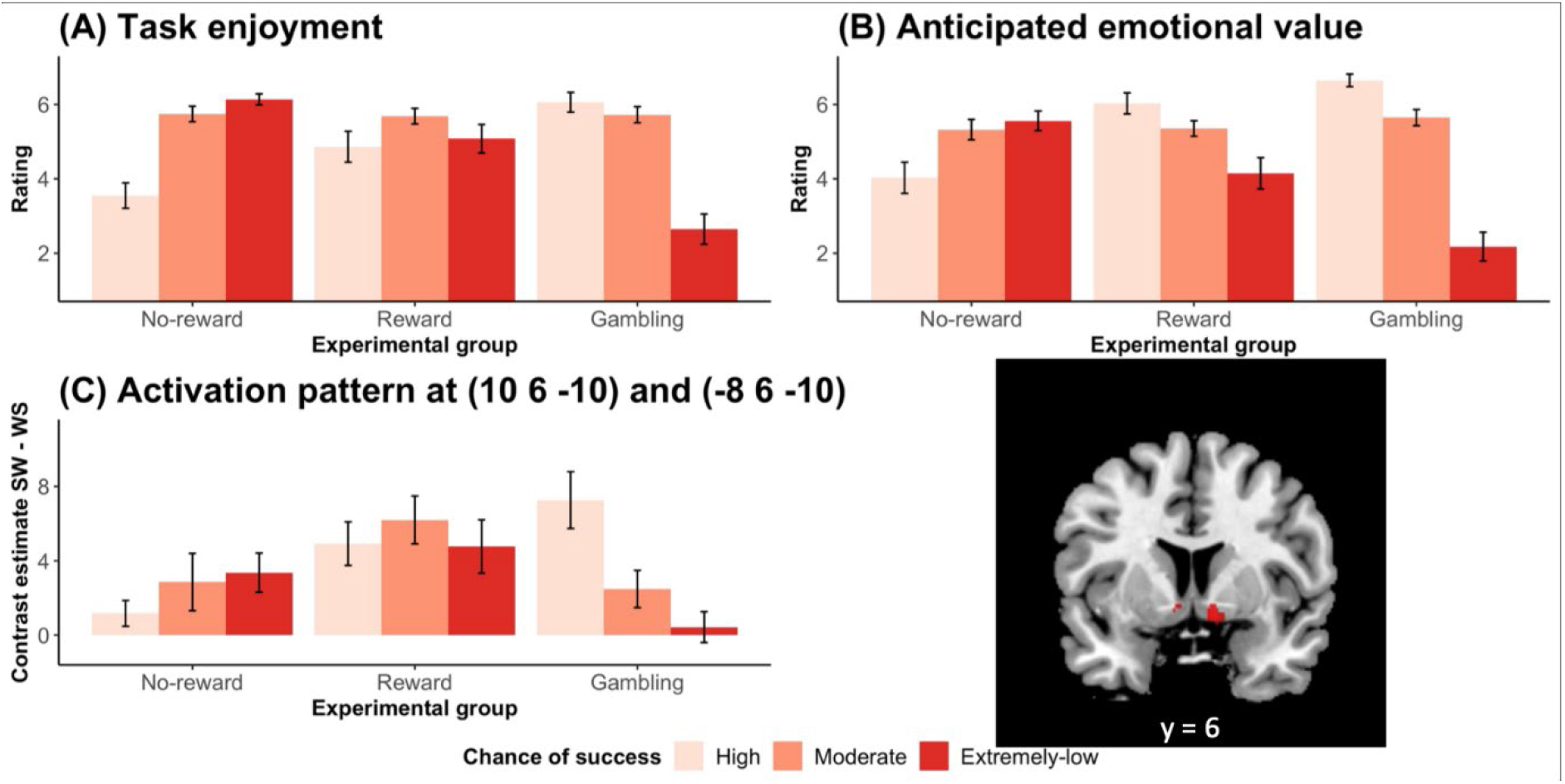
Pattern of results as a function of group (no-reward, reward, and gambling) and chance of success (high, moderately high, and extremely-low). **A**. Self-reported task enjoyment increased as the chance of success *decreased* in the no-reward group, whereas the pattern was opposite in the gambling group. Reward group is less clear, and none of the trend effects were significant. **B**. Self-reported anticipated emotional value for the task cue showed a similar pattern except that the anticipated emotional value monotonically decreased as the chance of success decreased in the reward group. **C**. The bilateral ventral striatum/ventral pallidum activation (right figure, *Ps* <.05, voxel-level FWE corrected) mirrored the pattern observed in the self-reported task enjoyment.

To confirm the pattern, we conducted a post-hoc trend analysis, examining the polynomial effects as a function of chance of success for each group. The analysis mostly supported our observation. In the no reward group, there was a significant linear trend of chance of success in the no reward group, *b* = − 2.59, *t* (32) – 7.32, *p* < .01, whereas this effect was qualified a negative quadratic effect, *b* = −1.80, *t* (32) = −2.95, *p* <.01. These results suggest that, consistent with our observation, there was an overall monotonic *increase* in task enjoyment as the chance of success decreased, although this increase was relatively smaller when the chance of success became extremely low perhaps due to a potential ceiling effect. In contrast, in the gambling group, there was a significant opposite linear trend, *b* = 3.41, *t* (32) = 7.33, *p* < .01, whereas this effect was somewhat compromised by a significant negative quadratic effect, *b* = −2.75, *t* (32) = −3.41, *p* < .01. These results suggest that, consistent with our observation, there was an overall monotonic *decrease* in task enjoyment as the chance of success decreased, although this decrease was relatively smaller when the chance of success changed from high to moderate. In the reward condition, nether the linear effect, *b* = −0.22, *t* (32) = −0.41, nor the quadratic effect, *b* = −1.43, *t* (32) = −1.59, was statistically significant.

For an exploratory purpose, we also assessed participants’ anticipated emotional value in response to the task cue for each of the three stopwatches (see Method). A similar ANOVA on the ratings of anticipated emotional value toward the task cue also showed a main effect of chance of success, *F*(2,96) = 23.87, *p* < .001, generalized *η*^2^ = 0.26, which was again qualified by an interaction effect, *F*(4,96) =23.39, *P* < .001, generalized *η^2^* = 0.41. The main effect of group was not statistically significant, *F*(2, 96) = 1.15. Interestingly and rather unexpectedly, the pattern was somewhat different from task enjoyment (Figure 2B). Participants in the no-reward and gamble groups followed the same trends as observed in the ratings for task enjoyment, while those in the reward group showed a *decreasing* trend as chance of success decreased: when the task became difficult (i.e. chance of success decreased), participants reported reduced anticipated emotional value for the task cue. Subsequent post-hoc trend analysis confirmed the observation: In all of the groups, there was a significant linear effect in a direction consistent with the observation, *b* = −1.53, *t* (32) = 3.05, *p* < .01; *b* = 1.88, *t* (32) = 4.15, *p* < .01; *b* = 4.47, *t* (32) = 11.76, *p* < .01; for the no-reward group, reward group, and gambling group, respectively. The quadratic effect was significant only for the gambling group, *b* = −2.47, *t* (32) = 3.75, *p* < .01, which seems to reflect the larger gap between the conditions of moderate chance of success and extremely-low chance of success. The other two groups did not show a significant quadratic effect, *b* = − 1.06, *t* (32) = −1.22 for the no-reward group; *b* = −0.53, *t* (32) = −0.67 for the reward group.

We also examined participants’ task performance. Although we manipulated performance feedback (see Methods; this ensures that participants’ subjective feelings of success were controlled across participants and groups), it was still possible to examine the objective performance regarding whether participants were able to press the button in the right timing, given that participants in the reward and no-reward groups were motivated to stop the stopwatch right at 5.00 seconds after the start of the stopwatch. We performed a 2 (chance of success; moderate chance vs. extremely-low chance) X 2 (group; no-reward vs. reward) ANOVA on the actual timing participants pressed a button. There was a significant main effect of the group, *F* (1, 32) = 4.79, *P* < 0.05, generalized *η^2^* = 0.06 suggesting that participants in the reward group stopped the stopwatch relatively earlier and less accurately (*M* = 4.919sec, *SD* = 0.065) than those in the no-reward group (*M* = 4.945sec, *SD* = 0.042 in the no-reward group). These results are consistent with the incentive-dependent performance decrements reported in previous research (Mobbs et al., 2009); pressures arising from performance-based extrinsic rewards may have impaired task performance. The main effect of chance of success and the interaction effect were not statistically significant (*Ps* > .05).

### Main GLM analysis of fMRI data

Our primary analysis focused on the reward network in the brain, especially the striatum/pallidum. Thus, we performed a region of interest (ROI) analysis. The striatum/pallidum ROI mask included the bilateral caudate, putamen, and pallidum from the Automated Anatomical Labeling (AAL) atlas (Tzourio-Mazoyer et al., 2002). The 3 (group) x 3 (chance of success) mixed ANOVA in our anatomical ROI revealed a significant interaction between group and chance of success in the bilateral ventral striatum extending into the ventral pallidum (VS/VP; Figure 2C; *Ps* < .05, voxel-level FWE corrected; *k* = 22 and 26 for the left and the right VS/VP, respectively). Confirming our hypothesis, the pattern of the results was consistent with the self-reported task enjoyment: whereas participants in the gambling group showed *decreased* activation in the VS/VP for task cues predicting lower-chance task of success, participants in the no-reward group showed *increased* activation for task cues predicting lower-chance of success. Participants in the reward group, on the other hand, showed a weaker effect, although the shape conforms to that of self-reported task enjoyment (i.e. an inverted-U shape). We also conducted a post-hoc trend analysis to confirm the pattern within the ROI (i.e., decreasing effect in gambling group, increasing effect in no-reward group, and inverted-U relationship in the reward group) and the analysis confirmed this pattern in the VS/VP (*P* < .05, voxel-level FWE corrected).

To demonstrate that these results do not simply reflect the general activation pattern in the entire brain, we also conducted a whole brain 3 × 3 ANOVA to explore other brain areas showing a similar pattern. The full results are reported in Table S1, and we also plotted the patten of activation across groups and conditions in Figure S1. Figure S1 also includes the exact location of the brain areas that showed a significant interaction effect. Both left and right VS/VP clusters survived with this stringent threshold, and the only other areas that survived were clusters in the visual areas. Activation patterns in the visual areas were generally similar with that of the VS/VP (Figure S1).

### Functional Connectivity Analysis

One interesting observation from the previous analyses is that, whereas participants in the no-reward group exhibited the highest task enjoyment (or the highest VS/VP activation) with extremely-low chance of success, participants in the reward group showed (at least numerically) decreased task enjoyment (and the striatal activation) for the cue for extremely-low chance of success relative to the cue for moderate chance of success. One possible explanation is that, in the reward group, participants’ task enjoyment for the near impossible task was compromised by the fact that they are very unlikely to obtain monetary rewards (Figure 1B). In fact, in our exploratory analysis, anticipated emotional value for the task cue in the reward group was lowest when the cue indicates extremely-low chance of success, suggesting the possibility that task enjoyment was curtailed by the anticipated emotional value (Figure 2B).

To examine the neural mechanisms underlying the decreased task enjoyment under the extremely-low chance of success in the reward group, we conducted a gPPI functional connectivity analysis (McLaren et al., 2012), which directly compared the functional connectivity in the extremely-low chance condition between the no-reward group and the reward group. The results showed that the activation in the left VS/VP (one of the seed regions) was more correlated with the activation in several brain areas in the reward group relative to the no-reward group. These areas include the left middle frontal gyrus, left inferior frontal gyrus, the right cerebellum, and left putamen/caudate (just outside of the ROI). Figure 3 shows these activated areas (all *P* < .05, cluster-level FWE corrected), and Table S2 shows peak coordinates. These results suggest that the left VS/VP and these areas are co-activated more strongly when participants in the reward group expected a task with little chance of success. The right VS/VP did not show such significant group differences in functional connectivity. Also, no-reward group did not show any significantly greater functional connectivity in comparison to reward group.

**Figure 3.**
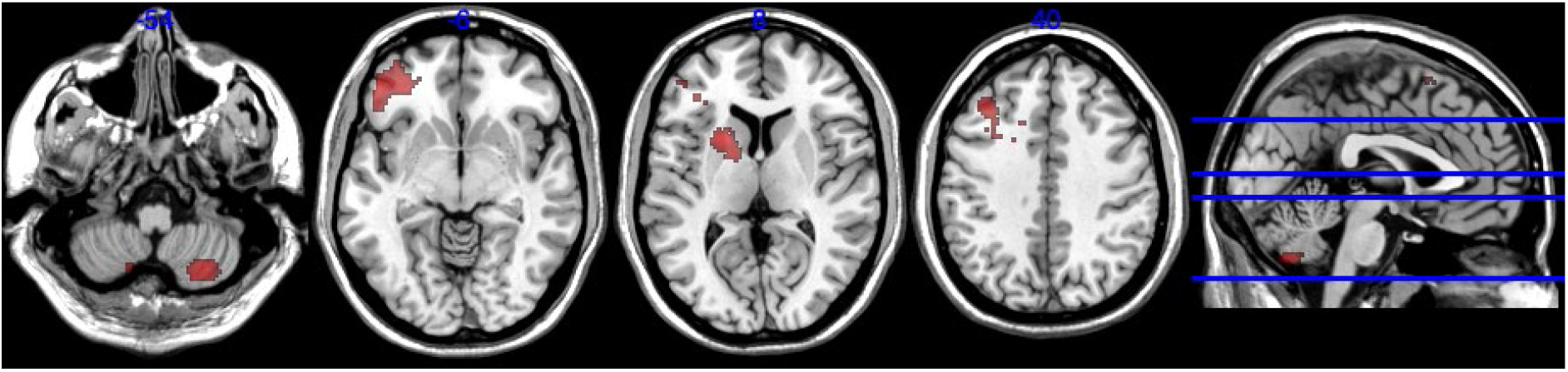
Results of generalized psychophysiological interactions analysis (gPPI; with the ventral striatum/ventral pallidum seed) comparing reward and no-reward groups for extremely-low chance of success. The results indicate that the activation in the left VS/VP was more correlated with the activation in the left middle frontal gyrus, left inferior frontal gyrus, left putamen/caudate, and the right cerebellum in the reward group, relative to the no-reward group (*P* < .05 with Bonferroni correction, cluster-level FWE-corrected).

## Discussion

The current study aimed to examine the factors influencing people’s motivation for a nearly impossible task and its neural correlates. Our results showed that, when the task success depended on participants’ skills (i.e. skill task) and did not promise any extrinsic rewards, participants’ overall motivation for the task (as assessed by task enjoyment) increased as the task became more difficult (i.e. lower chance of success), even to the extent that the task was almost impossible to achieve. However, when the task was skill-based but participants were promised performance-based monetary rewards, participants’ task enjoyment was hampered when the chance of success was extremely low; there was no clear relationship between the chance of success and task enjoyment. Also, when the task did not involve any skills and the outcome was determined in a purely probabilistic manner (i.e. luck task), the task enjoyment monotonically decreased as the chance of success decreased. Critically, VS/VP activation in response to the task cue basically mirrored these behavioral pattern of task enjoyment. Thus, the present research suggests that, both at the behavioral and neural levels, people are often intrinsically motivated to challenge a nearly impossible task but this happens mainly when the task requires certain skills and extrinsic incentives are not available.

Our findings pose some challenge to the normative economic and decision-making theories, because such a nearly impossible task should have little utility value given the extremely low expected success probability (Loewenstein, 1999). In fact, in the current experiment, participants never succeeded in the task of extremely-low chance of success. Using skill-based tasks, some previous neuroimaging studies have examined the effects of task difficulty by manipulating task demands (Dobryakova et al., 2017; Krebs et al., 2009; Kurniawan et al., 2010; Skvortsova et al., 2014; Ulrich et al., 2014). In these studies, however, participants in high demand task conditions exhibited decent task performance, making it difficult to disentangle the pure challenge-oriented motivation from the expected positive outcomes. In addition, most of the previous studies introduced extrinsic rewards in order to motivate participants toward an effortful task (except for a few studies; Boehler et al., 2011; Krebs et al., 2009). As our findings suggest, however, the presence of performance-based extrinsic rewards would make the interpretation of the results difficult, because such rewards provide external incentive to engage in a task with higher chance of success, which could be in direct conflict with people’s intrinsic inclination to work on a task of low chance of success.

Some existing theories have attempted to explain people’s challenge seeking behavior. For example, in the field of robotics and computational modelling, Oudeyer and his colleagues (Baranes, 2014; Oudeyer et al., 2013) argued that, when there are no explicit extrinsic rewards, agents may use learning progress as intrinsic rewards to motivate behavior. A recent empirical study confirmed that people are often motivated to work on a difficult task when they can expect learning progress (Ten et al., 2021). In the literature of psychology, researchers indicated that people seek (moderate) challenge because this provides people with feelings of competence and control (Csikszentmihalyi, 1990; Deci & Ryan, 1985), feelings of task achievement and pride (McClelland et al., 1976; Sedikides & Strube, 1997; see also Murayama et al., 2019), self-definition (Gendolla & Richter, 2010), and opportunity to reduce uncertainty to gauge one’s own ability (Trope, 1979). None of these theories, however, provide a viable explanation of why people are motivated for a nearly impossible task, because these theories consider occasional experiences of success as a critical source of motivation. One potential explanation for the observed motivation for the nearly impossible task is that participants were motivated to see how close they could be to succeeding in the task. In other words, in the absence of extrinsic rewards, participants might have gauged their achievement and progress against these internally generated sub-goals, rather than the externally-defined task success. In fact, in the current task, participants were able to see how close their button press was to the success. Future research should examine the exact psychological and neural mechanisms underlying the motivation for nearly impossible tasks.

This point also highlights the importance of distinguishing between skill tasks and luck tasks in decision-making and neuroscience research (Vostroknutov et al., 2012). A luck task (e.g. number guessing task) is a popular and predominant paradigm in decision science/neuroscience to examine how people or animals learn expected reward value from a task and make a decision. However, as noted above, people and animals also have the capacity to acquire rewarding value from the feeling of competence, acquiring knowledge, and achieving self-set goals (Gottlieb et al., 2013; Murayama, 2022; White, 1959). This means that more complex decision-making mechanisms might be operative with a task that entails some sort of skill learning or knowledge acquisition (Aarts et al., 2014; Hotaling et al., 2021; Knutson et al., 2005; Roiser et al., 2010). This aspect of the decision-making process is still underexamined in the literature, but worth attention for future research.

We observed an interesting dissociation between the task enjoyment and anticipated emotional value in response to the cue. The dissociation was specifically observed for an easy (high chance of success) task in the reward group: Whereas task enjoyment was relatively low (or not different) in this group, anticipated emotional value to the cue was the highest in comparison to other difficulty levels. These results indicate that, when the cue was presented, participants’ attention was primarily paid to the anticipated extrinsic rewards, not to the boresome experiences that they would be likely to feel in the actual task. This is consistent with the idea that people tend to underestimate the potential enjoyment/boredom of a task before working on it (Hatano et al., 2022; Kuratomi et al., 2018).

As discussed in the introduction, the reward network (including VS/VP) has been implicated in processing the expected reward value (or prediction error of a rewarding value, which is often confounded with the expected reward value). The current study provides some interesting insight into the role of the reward network in processing task value. Specifically, although the VS/VP activation patterns mirrored self-reported motivation, the parallel results were found with task enjoyment (i.e. positive emotional engagement in the task itself), rather than the emotional value toward the task cue (i.e. positive emotional feelings they experience on seeing the task cue). This is noteworthy, because we examined the brain activation in response to the task cue; nevertheless, VS/VP actually mirrored a positive emotional experience during the task, not the experience for the task cue. One possible interpretation is that activation of the reward network in this task might have reflected motivational relevance/mental preparation for an upcoming task (e.g., Carter, MacInnes, Huettel, & Adcock, 2009; Krebs, Boehler, Egner, & Woldorff, 2011). More specifically, it is possible that the striatum in the current experiment proactively integrated various sources of (both intrinsic and extrinsic) rewarding values based on the context in which the stimulus is presented (i.e. monetary reward value expected from the cue, emotional value of working on a challenging task, etc.), deciding the amount of future effort that should be allocated to the task. The idea that the striatum is related to the integrated net value is not new (e.g., Peters & Büchel, 2010). However, these arguments have been mainly limited to decision making that involves multiple attributes with concrete rewarding attributes (e.g., Magrabi et al., 2022), and little attention has been paid to the situation in which the reward context is manipulated (i.e. there are both intrinsic and extrinsic sources of rewarding values). Future study should further examine the role of the striatum in a skill-based task when the context of reward is explicitly manipulated.

Our functional connectivity analysis suggests that, when participants in the reward group expected a near impossible task, there is increased functional connectivity between the (left) VS/VP and some cortical areas ---the middle frontal gyrus and inferior frontal gyrus. In several meta-analyses, these cortical areas have been repeatedly implicated in the reappraisal of affective stimuli or events in the context of emotion regulation (Buhle et al., 2014; Frank et al., 2014; Picó-Pérez et al., 2019). Therefore, one possible interpretation is that working on a challenging task when there is performance-contingent rewards needs some cognitive reframing of the task, as there is direct cognitive conflict between people’s natural inclination to engage in a challenging task and people’s motivation to gain monetary rewards, as people are unlikely to succeed in a challenging task. Another interpretation is that this pattern may reflect the differences in cognitive control. Research has repeatedly indicated that the connectivity between the striatum and frontal cortex (i.e. frontal-striatal circuits) reflects cognitive control or executive functioning (e.g., Frank, 2011; Heyder et al., 2004; Westbrook et al., 2021). Therefore, the results may suggest that participants in the reward group decided to exert stronger cognitive control to focus on the task (at the sacrifice of task enjoyment), aiming to achieve the reward even while knowing that there is virtually no possibility. In any case, the current findings suggest the potential difficulty in interpreting the results when performance-based incentives are provided, because performance-based incentives could contradict with people’s tendency to engage in a challenging task.

The current study demonstrated that people have the inherent capability to motivate themselves for a nearly impossible task without extrinsic rewards, but failure is a major source of psychological dis-functioning. This is because failure in real life involves a multitude of factors that boost the negative implication of failure, such as social comparison, loss of future opportunities, and stigmatization. Previous studies have identified a number psychological underpinnings that are related to resilience and perseverance to failure (Duckworth et al., 2007; Oshio et al., 2018). Our work should be extended to accommodate such real-life factors, to better understand the potential and limitations of our intrinsic motivation to challenge very difficult tasks.

## Declarations

### Ethics approval

Study protocol was approved by the Ethics Committee of Tamagawa University.

### Consent of participate

Written, informed consent was obtained from participants

### Availability of data and materials

Unthresholded statistical maps are uploaded to NeuroVault (https://neurovault.org/collections/12642/).

### Code availability

Behavioral data and analysis code (including neuroimaging analysis) are uploaded to GitHub (https://github.com/stefaniemeliss/DME).

### Author contributions

K.Murayama., M.S., M.M., and K.Matsumoto conceived and designed research; M.M. programmed the task. K.Murayama, M.S., Y.Y., K.Matsumori, A.S., M.M., and K.Matsumoto conducted the experiment; K.Murayama, M.S., and S.M. analyzed data; K.Murayama, M.S., S.M. wrote the paper. All authors gave critical comments to the manuscript.

### Open Practices Statement

Unthresholded statistical maps are uploaded to NeuroVault (https://neurovault.org/collections/12642/). Behavioral data and analysis code (including neuroimaging analysis) is uploaded to GitHub (https://github.com/stefaniemeliss/DME). Raw imaging data are available on request. The reported experiment was not preregistered.

---

**Table S1.**
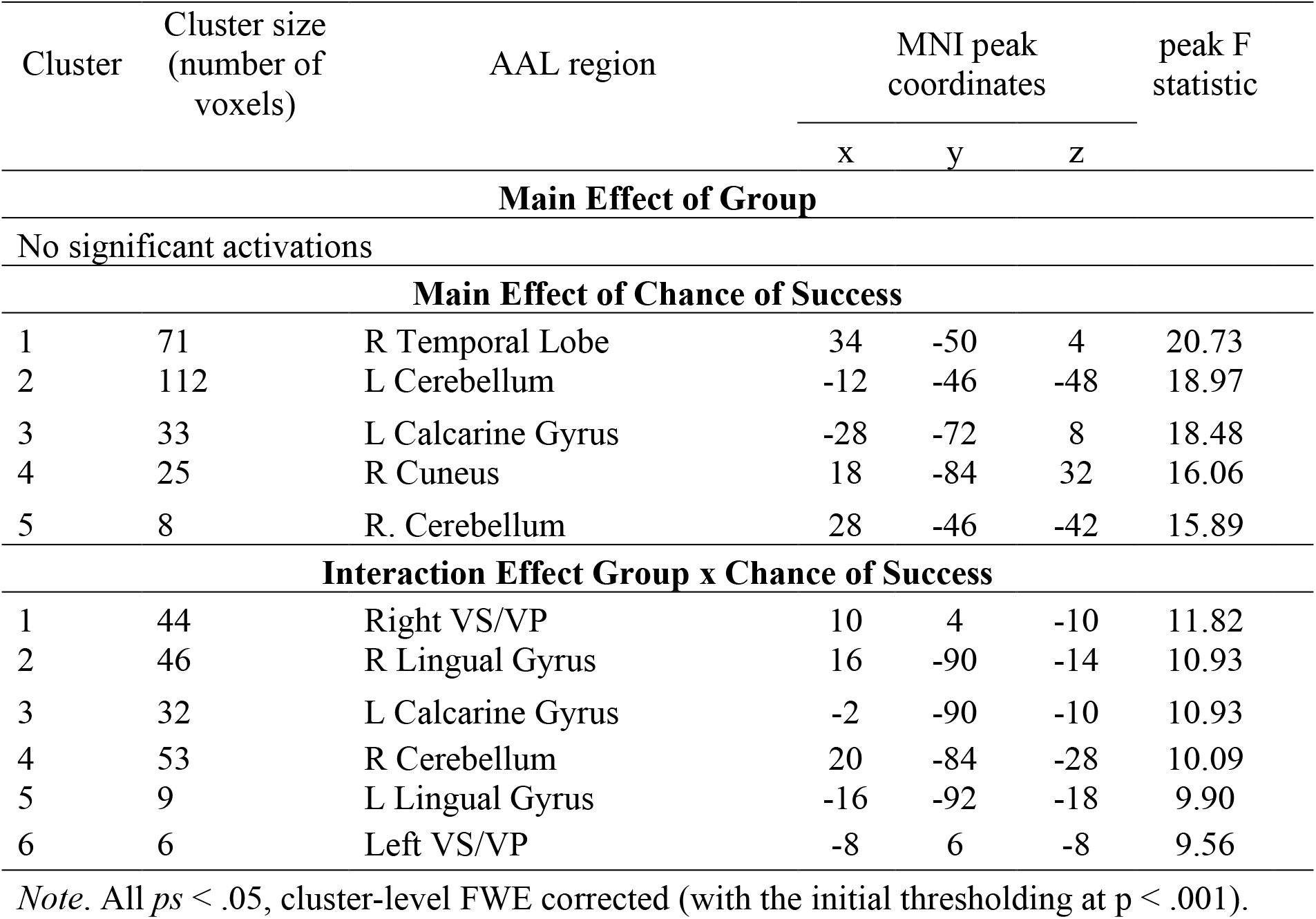
Results of 3 × 3 whole-brain ANOVA

| Cluster | Cluster size<br>(number of<br>voxels) | AAL region | MNI peak<br>coordinates |  |  | peak F<br>statistic |
| --- | --- | --- | --- | --- | --- | --- |
|  |  |  | x | y | z |  |
| Main Effect of Group |  |  |  |  |  |  |
| No significant activations |  |  |  |  |  |  |
| Main Effect of Chance of Success |  |  |  |  |  |  |
| 1 | 71 | R Temporal Lobe | 34 | -50 | 4 | 20.73 |
| 2 | 112 | L Cerebellum | -12 | -46 | -48 | 18.97 |
| 3 | 33 | L Calcarine Gyrus | -28 | -72 | 8 | 18.48 |
| 4 | 25 | R Cuneus | 18 | -84 | 32 | 16.06 |
| 5 | 8 | R. Cerebellum | 28 | -46 | -42 | 15.89 |
| Interaction Effect Group x Chance of Success |  |  |  |  |  |  |
| 1 | 44 | Right VS/VP | 10 | 4 | -10 | 11.82 |
| 2 | 46 | R Lingual Gyrus | 16 | -90 | -14 | 10.93 |
| 3 | 32 | L Calcarine Gyrus | -2 | -90 | -10 | 10.93 |
| 4 | 53 | R Cerebellum | 20 | -84 | -28 | 10.09 |
| 5 | 9 | L Lingual Gyrus | -16 | -92 | -18 | 9.90 |
| 6 | 6 | Left VS/VP | -8 | 6 | -8 | 9.56 |

*Note.* All $ps < .05$ , cluster-level FWE corrected (with the initial thresholding at $p < .001$ ).

**Table S2.** Results of gPPI analysis with the ventral striatum/ventral pallidum seed, comparing reward and no-reward groups for extremely-low chance of success.

| Cluster | Cluster size<br>(number of<br>voxels) | AAL region | MNI peak<br>coordinates |  |  | peak t<br>statistic |
| --- | --- | --- | --- | --- | --- | --- |
|  |  |  | x | y | z |  |
| Seed: Right VS/VP, reward group > no-reward group |  |  |  |  |  |  |
| No significant activations |  |  |  |  |  |  |
| Seed: Right VS/VP, no-reward group > reward group |  |  |  |  |  |  |
| No significant activations |  |  |  |  |  |  |
| Seed: Left VS/VP, reward group > no-reward group |  |  |  |  |  |  |
| 1 | 267 | L Putamen | -22 | 12 | 8 | 6.04 |
| 2 | 564 | R Cerebellum | 30 | -64 | -54 | 5.47 |
| 3 | 588 | L Middle Frontal Gyrus | -36 | 30 | 40 | 5.41 |
|  |  | L Somatosensory Area | -28 | 14 | 32 |  |
| 4 | 828 | L Inferior Frontal Gyrus | -48 | 40 | -6 | 5.35 |
|  |  | L Middle Orbital Gyrus | -32 | 42 | -8 |  |
| Seed: Left VS/VP, no-reward group > reward group |  |  |  |  |  |  |
| No significant activations |  |  |  |  |  |  |

*Note.* All $ps < .05$ , cluster-level FWE corrected (with the initial thresholding at $p < .0025$ ).

**Figure S1.**
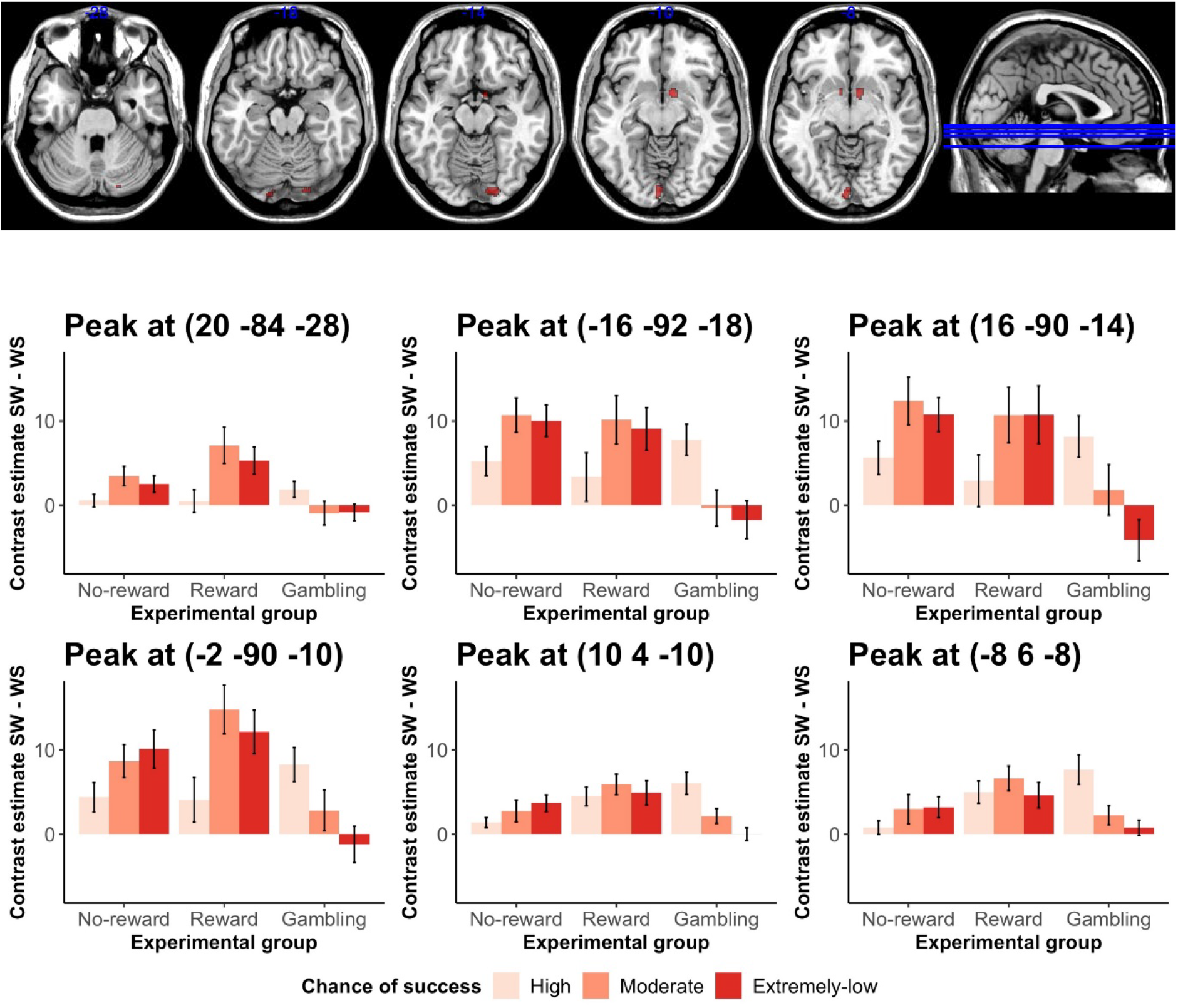
Activation pattern across conditions and groups for the peak voxels of the clusters that showed a significant 2-way interaction (with activation map to indicate the exact locations).

